# Reelin deficiency exacerbates cocaine-induced hyperlocomotion by enhancing neuronal activity in the dorsomedial striatum

**DOI:** 10.1101/2022.04.12.488105

**Authors:** Giordano de Guglielmo, Attilio Iemolo, Aisha Nur, Andrew Turner, Patricia Montilla-Perez, Francesca Telese

**Author notes:** <u>Correspondence:</u> Francesca Telese, Department of Medicine, University of California San Diego, La Jolla, CA, 92093, USA.

## Abstract

The *Reln* gene encodes for the extracellular glycoprotein Reelin, which regulates several brain functions from development to adulthood, including neuronal migration, dendritic growth and branching, and synapse formation and plasticity. Human studies have implicated Reelin signaling in several neurodevelopmental and psychiatric disorders. Mouse studies using the heterozygous Reeler (HR) mice have shown that reduced levels of *Reln* expression are associated with deficits in learning and memory and increased disinhibition. Although these traits are relevant to substance use disorders, the role of Reelin in cellular and behavioral responses to addictive drugs remains largely unknown. Here, we compared HR mice to wild-type (WT) littermate controls to investigate the contribution of Reelin signaling to the hyper-locomotor and rewarding effects of cocaine. After a single cocaine injection, HR mice showed enhanced cocaine-induced locomotor activity compared to WT controls. After repeated injections of cocaine, Reelin deficiency also led to increased cocaine-induced locomotor sensitization, which persisted after withdrawal. In contrast, Reelin deficiency did not affect the rewarding effects of cocaine measured in the conditioned place preference assay. The elevated cocaine-induced hyper-locomotion in HR mice resulted in increased *Fos* expression in the dorsal medial striatum (DMS) compared to WT. Lastly, we found that *Reln* was highly co-expressed with the *Drd1* gene, which encodes for the dopamine receptor D1, in the DMS.

These findings demonstrated that Reelin signaling contributes to the locomotory effects of cocaine and improved our understanding of the neurobiological mechanisms underlying the cellular and behavioral effects of cocaine.

## Introduction

Reelin is an extracellular glycoprotein expressed by Cajal Retzius cells during brain development and predominately by GABAergic interneurons in the postnatal brain ^1–3^. The canonical Reelin signaling pathway includes the binding of Reelin to lipoprotein membrane receptors ApoER2 and VLDR, the phosphorylation of the intracellular adaptor protein Dab1by two tyrosine kinases of the Src family, Src and Fyn, and the recruitment of singling molecules of the Crk family ^4–7^. During embryonic development, Reelin signaling is required for the proper migration of newborn neurons involved in the laminar formation of several brain structures, such as the cortex, hippocampus, and cerebellum ^2,8^. Mutant mice lacking *Reln* (Reeler mice) or other core components of the Reelin signaling show grossly inverted cortical layers due to defective neuronal positioning ^9^. In humans, *Reln* gene mutations are associated with autosomal recessive lissencephaly with cerebellar hypoplasia, characterized by gross brain malformations similar to those observed in the Reeler mice ^10^.

In the postnatal period, Reelin signaling is implicated in several brain functions linked to cognition, including dendritic spine formation, excitatory glutamatergic synaptic activity, and activity-regulated transcription ^11–19^. The Heterozygous Reeler (HR) mice express reduced levels of Reelin and do not exhibit abnormal neuronal positioning during development ^20^. In contrast, they show reduced dendritic spine density ^21^ and several behavioral abnormalities, including impaired learning and memory, decreased behavioral inhibition, and increased impulsivity ^20,22–25^. Therefore, HR mice have been proposed as a model for studying neurological disorders ^20^. A link between Reelin signaling and psychiatric diseases is strongly supported by human genetic studies or postmortem brain studies showing that perturbation of Reelin signaling is associated with schizophrenia, bipolar disorder, autism, epilepsy, and Alzheimer’s disease ^26–28^.

Despite the evidence implicating Reelin signaling in several psychiatric disorders, including behavioral traits relevant to addiction, the influence of Reelin on the cellular and behavioral effects of addictive drugs remains largely unknown.

Here, we used the HR mice to assess the influence of reduced levels of Reelin on the hyperlocomotion and reward effects of cocaine, a psychostimulant that leads to robust cellular and behavioral changes in mice. We measured the effects of acute or repeated injection of cocaine on locomotor activity in HR and WT littermate controls. We used a conditioned place preference (CPP) assay to evaluate the effects of Reelin deficiency on the rewarding effects of cocaine. In addition to behavioral tests, we mapped the regional expression of the cocaine-induced *Fos* expression in the dorsomedial striatum (DMS) and the nucleus accumbens shell (NAc Shell) in HR and WT mice. Finally, we performed RNA fluorescent in situ hybridization to examine *Reln* expression in specific cell types of the DMS.

## Materials and Methods

### Subjects

All experimental procedures were approved by the institutional animal care and use committee at the University of California, San Diego. Mice were housed (3–4 per cage) under a 12h light/12h dark cycle and provided with food and water ad libitum. HR mice were bred in house using the B6C3Fe a/a-Relnrl/J line (The Jackson Laboratory, #000235)^9^. Mice used for the locomotor activity and CPP were transferred at the TSRI Behavioral Core.

### Genotyping

Genotyping was carried out by PCR using genomic DNA extracted from tail clips. PCR was performed using the DreamTaq Hot Start Green PCR Master Mix (Thermo Scientific, Cat. no. K1082), and the following primers: a forward primer common to both alleles 5’TAATCTGTCCTCACTCTGCC 3’, a reverse wildtype specific primer 5’ACAGTTGACATACCTTAATC3’ and a reverse mutant specific primer 5’TGCATTAATGTGCAGTGTTG3’. The expected sizes for the amplicons are 280bp for the WT and 280bp + 380bp for the HET.

### Drugs

Cocaine HCl (National Institute on Drug Abuse, Bethesda, MD, USA) was dissolved in 0.9% saline (Hospira, Lake Forest, IL, USA) at various concentrations (5,10,20, and 40 mg/kg) and injected intra-peritoneally (IP).

### Locomotor activity chambers

Locomotor activity was measured in polycarbonate cages (42 x 22 x 20 cm) placed into frames (25.5 x 47 cm) mounted with two levels of photocell beams at 2 and 7 cm above the bottom of the cage (San Diego Instruments, San Diego, CA). These two sets of beams allowed for recording both horizontal (locomotion) and vertical (rearing) behavior. A thin layer of bedding material was applied to the bottom of the cage.

### Cocaine-induced locomotor activity in HR and WT mice

The mice (20 HR and 21 WT, half males and half females) were tested immediately following IP administration of 0.9% saline solution (0.01ml/g body weight) or 10 mg/kg cocaine HCl with locomotor activity recorded for 15 min (in 1 min bins).

### Dose-response of cocaine-induced locomotor activity in HR and WT mice

The mice (26 HR and 23 WT, half males and half females) were first tested immediately following IP administration of 0.9% saline solution (0.01ml/g body weight) with locomotor activity recorded for 15 min (in 1 min bins) to acclimate them to the test. Starting 3 days later, mice received 0.0 (saline), 5, 10, 20, and 40 mg/kg of cocaine HCl in a counterbalanced order, one dose every 5 days, and were tested in the locomotor activity cages for 15 min. In this way, a within-subjects dose-response curve was generated for each group.

### Cocaine-induced locomotor sensitization in HR and WT mice

All mice (10 HR and 8 WT were first tested immediately following IP administration of 0.9% saline solution (0.01ml/g body weight) with locomotor activity recorded for 15 min (in 1 min bins) on two consecutive days (Days 1 & 2) to acclimate them to the test. Mice then received 10 mg/kg cocaine on Days 3, 5, 7, 9, and 11 and were tested in the locomotor activity cages for 15 min. Intermittent cocaine exposures were used to examine the progression of the behavioral sensitization ^29–31^. Finally, on Day 12, all mice received saline before an identical locomotor test. One week later, all mice received 10 mg/kg of cocaine to examine sensitivity to the same dose used in the sensitization test.

### Cocaine conditioned place preference (CPP) in HR and WT mice

Place conditioning involves pairing a distinct environmental context (i.e., floor type) with a motivationally significant event (i.e., cocaine injection) ^32^. Rectangular Plexiglas black matte boxes (length 42 cm, width 22 cm, height 30 cm) divided by central partitions into two chambers of equal size (22×22×30 cm) were used. Distinctive tactile stimuli were provided in the two compartments of the apparatus. One chamber had no additional flooring (i.e. was smooth), and the other had lightly textured milky-colored flooring. During pre-conditioning and testing sessions, an aperture (4×4 cm) in the central partition allowed the animals to enter both sides of the apparatus. All testing occurred during the dark cycle under red light and was analyzed from video files using Noldus Ethovision software.

42 mice were used for this experiment, 20 HR and 22 WT, half males and half females. The experiment consisted of three phases: pre-conditioning, conditioning, and testing in the following sequence. Day 1: Pre-conditioning phase with access to both compartments, Day 2-7: Conditioning phase - Drug or saline administration followed by immediate confinement in one compartment of the place conditioning apparatus, and Day 8: Test with access to both compartments in the drug-free state. For the pre-conditioning phase, each animal was placed in one compartment of the apparatus and allowed to explore the entire apparatus for 30 min. The time spent in each of the two compartments was measured. Mice showing unbiased exploration of the 2 sides of the apparatus (between 45 and 55% time spent on each side) were randomly assigned a chamber in which to receive cocaine. Mice showing biased exploration of either side were given cocaine in the least preferred compartment. Bias was observed in half of the mice tested and this was spread equally across each genotype and therefore did not confound the results. On the following 6 days, 30 min conditioning sessions were given in which animals were injected i.p. with either saline or 10 mg/kg cocaine and immediately confined to one side of the apparatus (alternating these treatments and sides each day). In this way, each mouse experienced 3 pairings of cocaine with one of the apparatus sides. On the day after the final conditioning trial, each mouse was allowed to explore the entire apparatus in a non-drugged state for 30 minutes. The time spent in each of the two compartments of the conditioned place preference apparatus was recorded. The time spent in the cocaine side was compared with the pre-conditioning session to examine preference development.

### Immunohistochemistry (IHC)

To examine Fos expression by IHC, we used 4 female WT and 4 female HET littermates. Mice were injected with 10mg/kg of cocaine. One hour after injection, mice were anesthetized with CO2 and fixed via transcardial perfusion with 4% paraformaldehyde (PFA) in phosphate-buffered saline (PBS). Brain tissues were processed at a cryostat, as previously described ^33^. 30-μm thick coronal slices were stained with a primary antibody recognizing Fos (1:1000, ABE457, EMD Millipore, RRID:AB_2631318) and a secondary anti-rabbit antibody (donkey anti-rabbit Alexa Fluor 488 (1:1000, #A21206, Thermo Fisher Scientific). Images were acquired with a fluorescent microscope (BZX800, Keyence Corporation, Osaka, Japan). Different brain areas were identified using a mouse brain atlas as a reference ^34^. For each region, we stained 4-5 sections from Bregma 0.5mm to 1.94mm, which span both areas. The number of Fos+ cells for each mouse was normalized to the size of the chosen area (mm2) and presented as mean Fos + per mm^2^± SEM. One outlier (WT-DMS) was removed from the analysis (identified after the Grubbs’ test with alpha set at 0.05).

### RNA fluorescent in situ hybridization (FISH)

To examine the expression of various transcripts in specific cell types of the DMS, we performed RNA FISH using the RNAscope Multiplex Fluorescent Reagent Kit v2 (ACD, #323100) and following the instructions for the ‘fixed-frozen tissue sample’ of the user manual (ACD, USM-323100). 30-μm thick coronal slices were obtained from 2 WT mice, as described above. Tissue sections corresponding to DMS (Bregma 0.86mm) from 2 WT mice were hybridized with a mix of three probes; Reelin (ACD, # 405981) + Drd1 (ACD, 406491-C2) + Drd2 (ACD, 406501-C3). We used DAPI as a nuclear stain. To assess both tissue RNA integrity and assay procedure, a separate group of sections was incubated with negative probes (data not shown). Images were acquired with the Keyence fluorescent microscope. The number of nuclei was counted manually and normalized to the size of the area selected (mm^2^).

### Statistical Analysis

The data are expressed as mean ± SEM. For comparisons between only two groups, *p* values were calculated using unpaired *t*-tests as described in the result section. Comparisons across more than two groups were made using one-way analysis of variance (ANOVA), and two- or three-way ANOVA was used when there was more than one independent variable. ANOVA was followed by Bonferroni’s multiple comparison test when appropriate. Differences were considered significant at *p* < 0.05. The standard error of the mean is indicated by error bars for each data group. All of these data were analyzed using Statistica 7 software.

## Results

### HR mice exhibit enhanced hyper-locomotory effects of cocaine compared to WT

We injected HR and WT mice with 10mg/kg of cocaine or saline and measured their locomotor activity for the first 15 minutes after injection. The three-way ANOVA with genotype (HR and WT) and dose (0.0 and 10 mg/kg) as between factors and the time as within factor indicated a significant genotype * dose interaction (F_1,555_=49.15; *p*<0.001), demonstrating that HR rats showed increased locomotor activity compared to WT after a single injection of 10 mg/kg of cocaine (Fig. 1A). This effect was confirmed when the data were presented as the average locomotor activity during the 15 minutes test recording (Fig. 1B). The two-way ANOVA of the mean locomotor activity with genotype (HR and WT) and dose (0.0 and 10 mg/kg) as between factors showed a significant genotype * dose interaction (F_1,37_=4.09; *p* = 0.05). Post hoc comparisons with Bonferroni’s test corrections demonstrated that cocaine significantly increased the mean locomotor activity compared to saline in both genotype groups (*p* < 0.001 vs. saline). This effect was significantly higher in the HR mice compared to WT (*p* < 0.01, Fig. 1B).

**Figure1.**
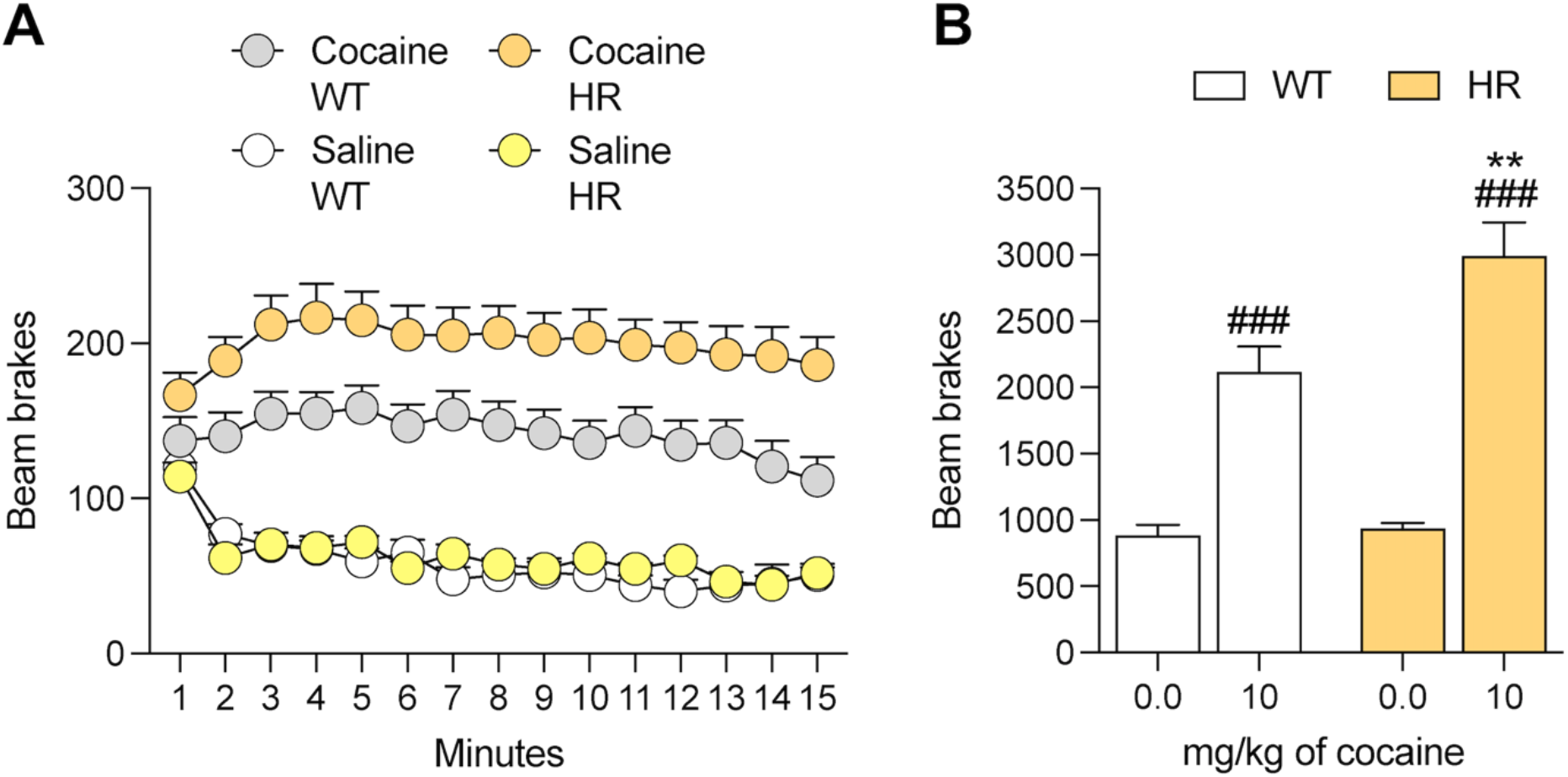
Locomotor activity in HR and WT mice after a single injection of cocaine. **A**) Number of beam brakes per minute during the 15 minutes test. **B**) Total number of beam brakes during the 15 minutes locomotor test. ** *p* < 0.01 vs WT; *###p* < 0.001 vs dose 0.0 (saline).

### Dose-response of cocaine-induced locomotor activation in HR and WT mice

We compared the locomotor activity of WT and HR mice after the injection of different doses of cocaine. The two-way ANOVA of the mean locomotor activity with genotype (HR and WT) and dose (0.0 and 10 mg/kg) as between factors showed a significant effect of the genotype (F_1,46_=9.82; *p* < 0.01), of the dose (F_4,46_=27.64; *p* < 0.0001), but no genotype * dose interaction, demonstrating that cocaine increases locomotor activity in HR mice independently from the dose used (Fig. 2A). We then analyzed the data by adding sex as a factor to examine the influence of sex on the increased stimulatory effects of cocaine observed in HR mice. The three-way ANOVA with genotype (HR and WT), cocaine dose and sex as factors did not show any sex * genotype (F_1,220_=13.39; *p* = 0.09) or sex * genotype * dose interactions (F_4,220_=0.084; *p* = 0.98), demonstrating that sex does not play a role in the increased stimulatory effects of cocaine in HR mice (Fig. 2B).

**Figure 2.**
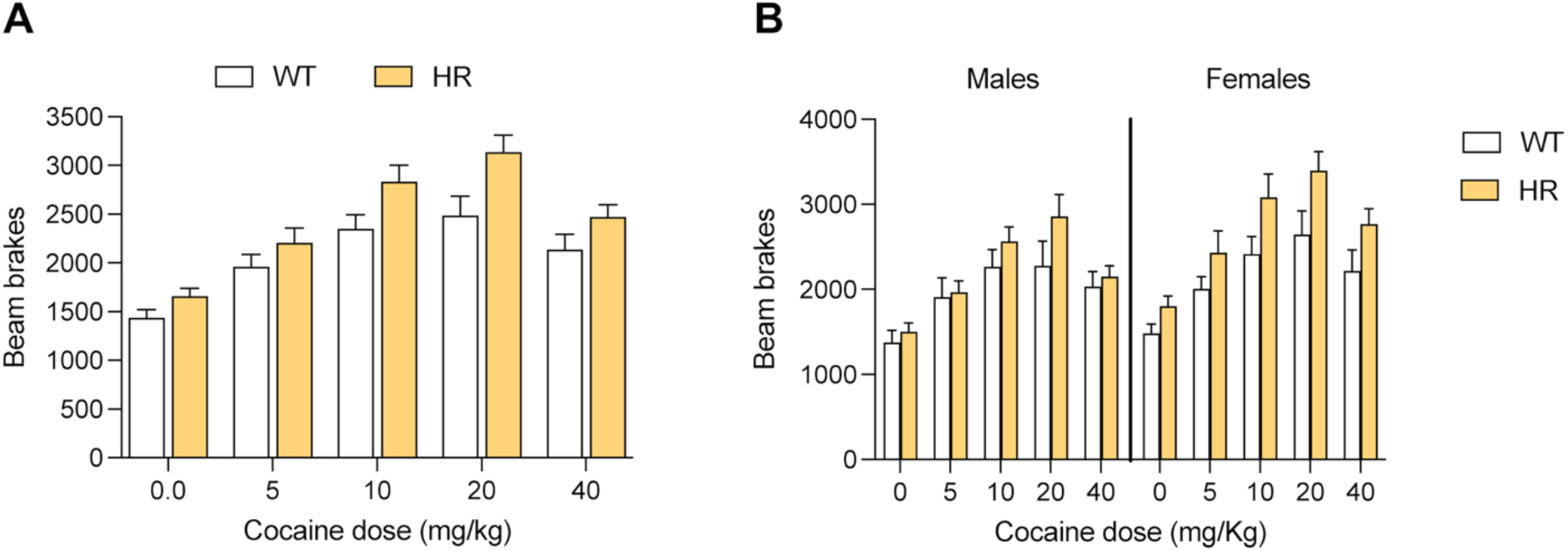
Dose-response for the effects of a single cocaine injection on locomotor activity in HR and WT mice. **A**) Dose-response for the total population. **B**) Dose-response in male and female mice.

### HR mice show increased cocaine-induced locomotor sensitization compared to WT

We explored the development of behavioral sensitization in WT and HR mice following repeated exposures to 10mg/kg of cocaine (Fig. 3A). No significant differences in locomotion were observed between the HR and WT mice in the first 2 days while receiving saline injections (Fig. 3B). This observation indicates that Reelin deficiency does not impact novelty-induced hyperlocomotion or habituation to the novel environment. Over the course of the experiment, HR mice injected with cocaine showed a dramatic increase in locomotor activity compared to WT, as demonstrated by the significant genotype effect on the “cocaine days” (F_1,16_=4.72; *p* < 0.05) in a Two-way ANOVA (Fig. 3B). There were no detectable differences in locomotion on day 12 when the animals were re-exposed to saline. A comparison of the locomotion values on day 19, when the mice received a new cocaine injection after one week of washout, showed that cocaine-treated HR mice presented significantly higher locomotion values than WT mice (t_17_ = 2.171; *p* < 0.05, Fig. 3B).

**Figure 3.**
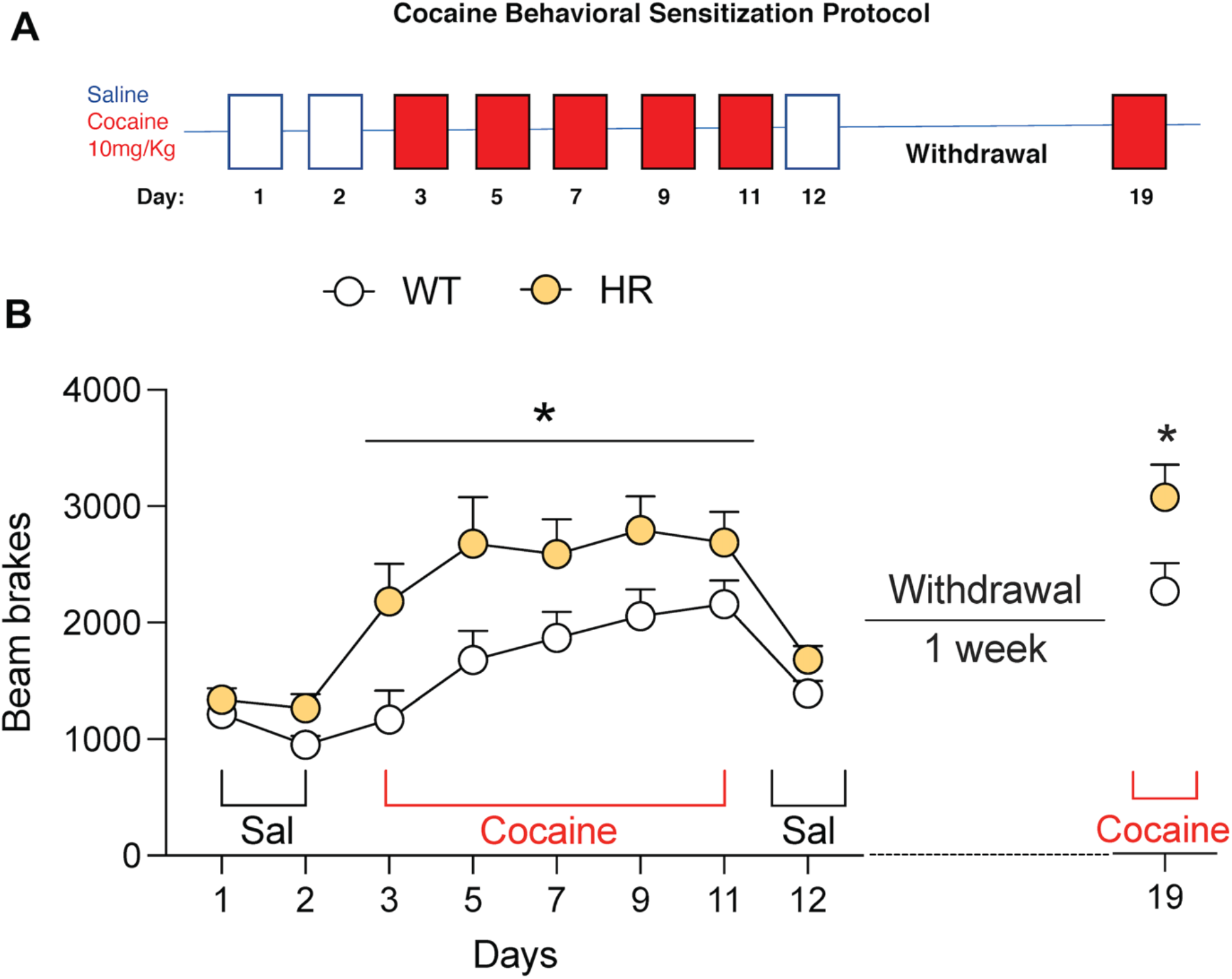
Reelin downregulation increases cocaine sensitization in HR mice. **A**) Schematic timeline of the experiment. **B**) Development of locomotor sensitization in HR and WT mice. * *p* < 0.05 vs WT

### Reelin deficiency does not influence the rewarding effects of cocaine

To investigate the impact of Reelin deficiency on the rewarding effects of cocaine, we compared WT and HR mice in the CPP assay to measure the preference of mice for a context associated with the cocaine reward. The preference is expressed as the time spent in the compartment associated with cocaine. The two-way ANOVA with genotype as between factor and preference (BSL vs. Test) as within factor showed a significant effect of preference (F_1,40_=129.2; *p* < 0.001, Fig. 4A), but not detectable genotype effect, demonstrating that cocaine induced a robust place preference in both HR and WT mice.

**Figure 4.**
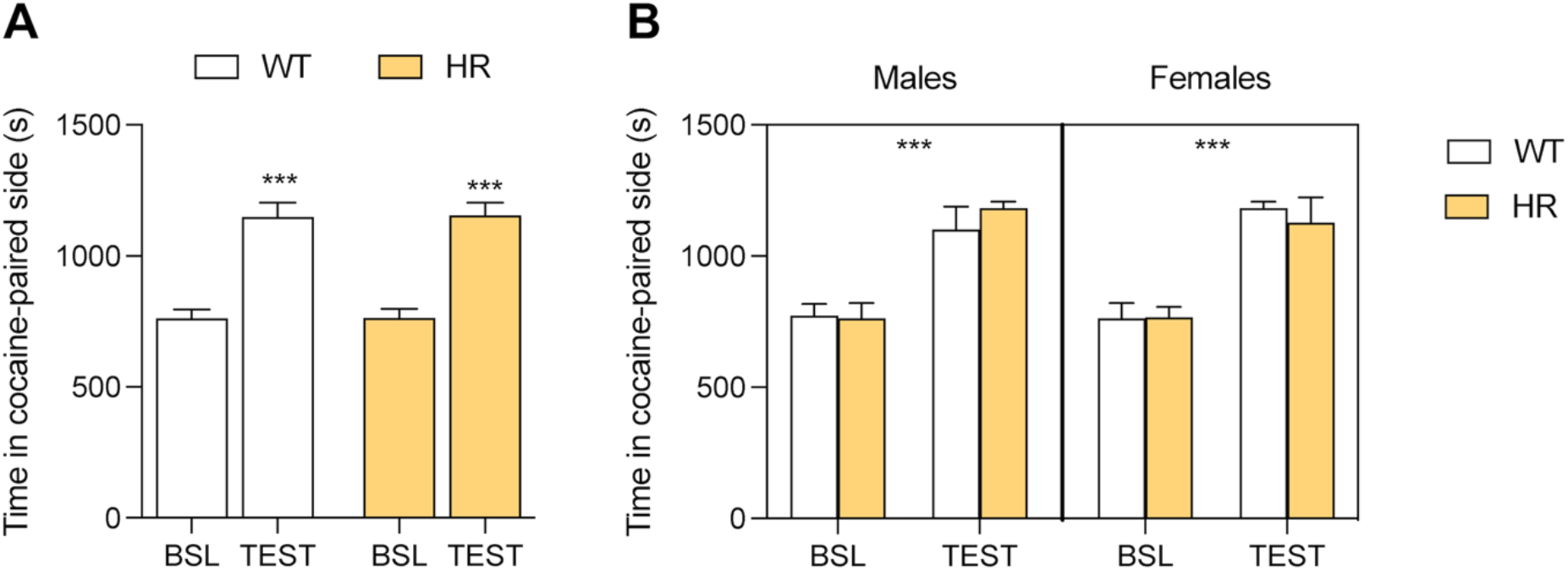
Reelin downregulation does not affect the rewarding properties of cocaine in a place conditioning paradigm. **A**) Time spent in the cocaine-paired side in HR and WT mice. **B**) Place conditioning results in male and female mice. ** *p* < 0.01 vs baseline (BSL).

When sex was added as a factor in the analysis, the three-way ANOVA showed a main effect of the preference (F_1,38_=150.1; *p* < 0.0001, Fig. 4B) without detectable sex effects, demonstrating that cocaine induced a robust place preference in both HR and WT mice, independently of the sex.

### Cocaine-induced hyper-locomotion in HR mice is associated with increased *Fos* expression in the dorsal medial striatum

To gain deeper insights into the neuronal ensembles underlying the increased locomotor activity enhanced by cocaine in HR mice, we mapped the regional expression of Fos protein by IHC in DMS and NAc shell, two regions implicated in the acute behavioral responses to cocaine. The two-way ANOVA with genotype and brain region as factors showed a significant genotype * brain region interaction (F_1,13_ =5.958;*p* < 0.05, Fig. 5). The posthoc comparisons with the Bonferroni’s correction demonstrated that, after a single injection of cocaine, HR mice showed increased levels of Fos positive cells in the DMS, but not in the NAc shell, compared to WT mice (adj *p*< 0.01, Fig. 5).

**Figure 5.**
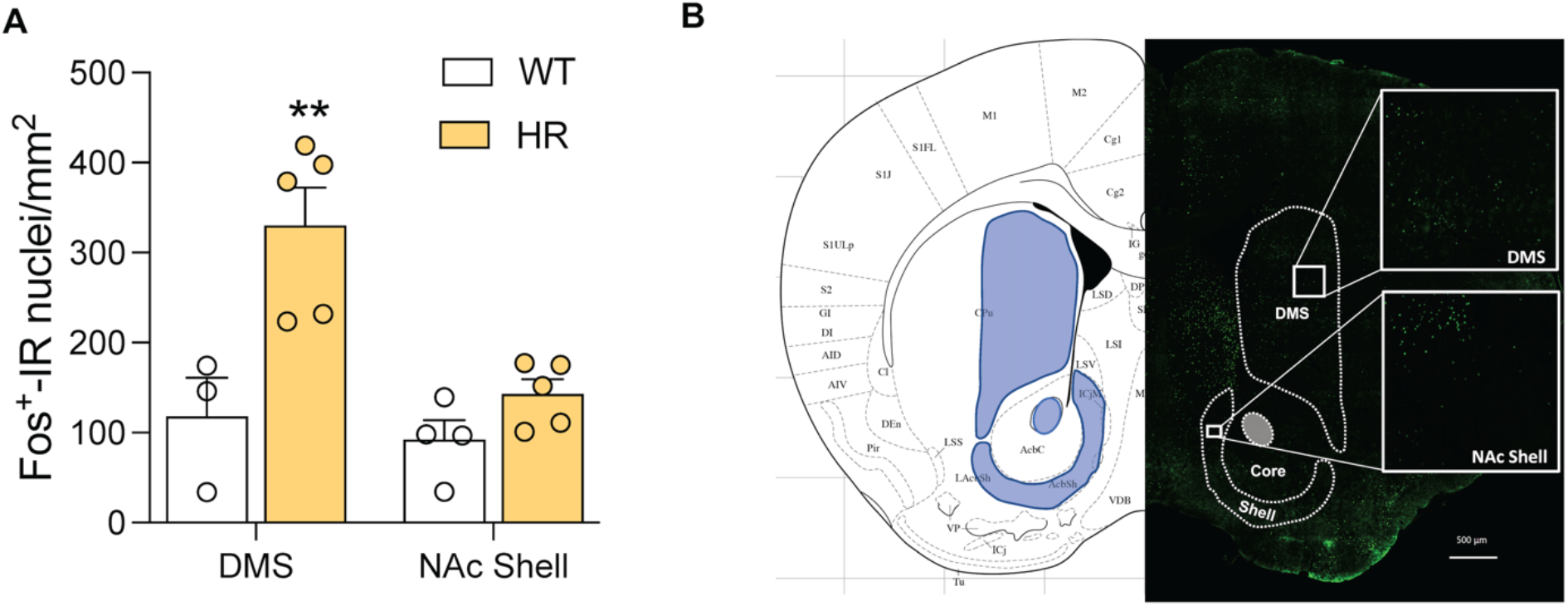
HR mice showed higher levels of Fos+ cells in the DMS compared to WT after a single injection of cocaine. **A**) Fos+ cells in the DMS and the NAc shell. **B**) Representative images of the immunostaining. ** *p* < 0.01 vs WT

### Reelin is highly coexpressed with Drd1 in the dorsal medial striatum

Acute responses to cocaine lead to increased dopamine levels in the ventral tegmental area and are mediated by neuronal populations expressing the dopamine receptors, Drd1 and Drd2, which are highly expressed in the medium spiny neurons (MSNs) of DMS. We analyzed *Reln* expression in the Drd1 and Drd2 subpopulations of the DMS by RNA FISH. While similar expression of *Drd1* and *Drd2* transcripts was detected in the DMS (Fig. 6A), we found increased co-expression of *Reln* transcript in Drd1+ compared to Drd2+ neurons, as demonstrated by the t-test (t_6_ = 2.486; *p* < 0.05, Fig 6B). When data were transformed in percentage, we found that 67% of the Drd1+ cells and 46% of the Drd2+ cells co-expressed *Reln* (t_6_ = 2.486; *p* < 0.05, Fig 6C).

**Figure 6.**
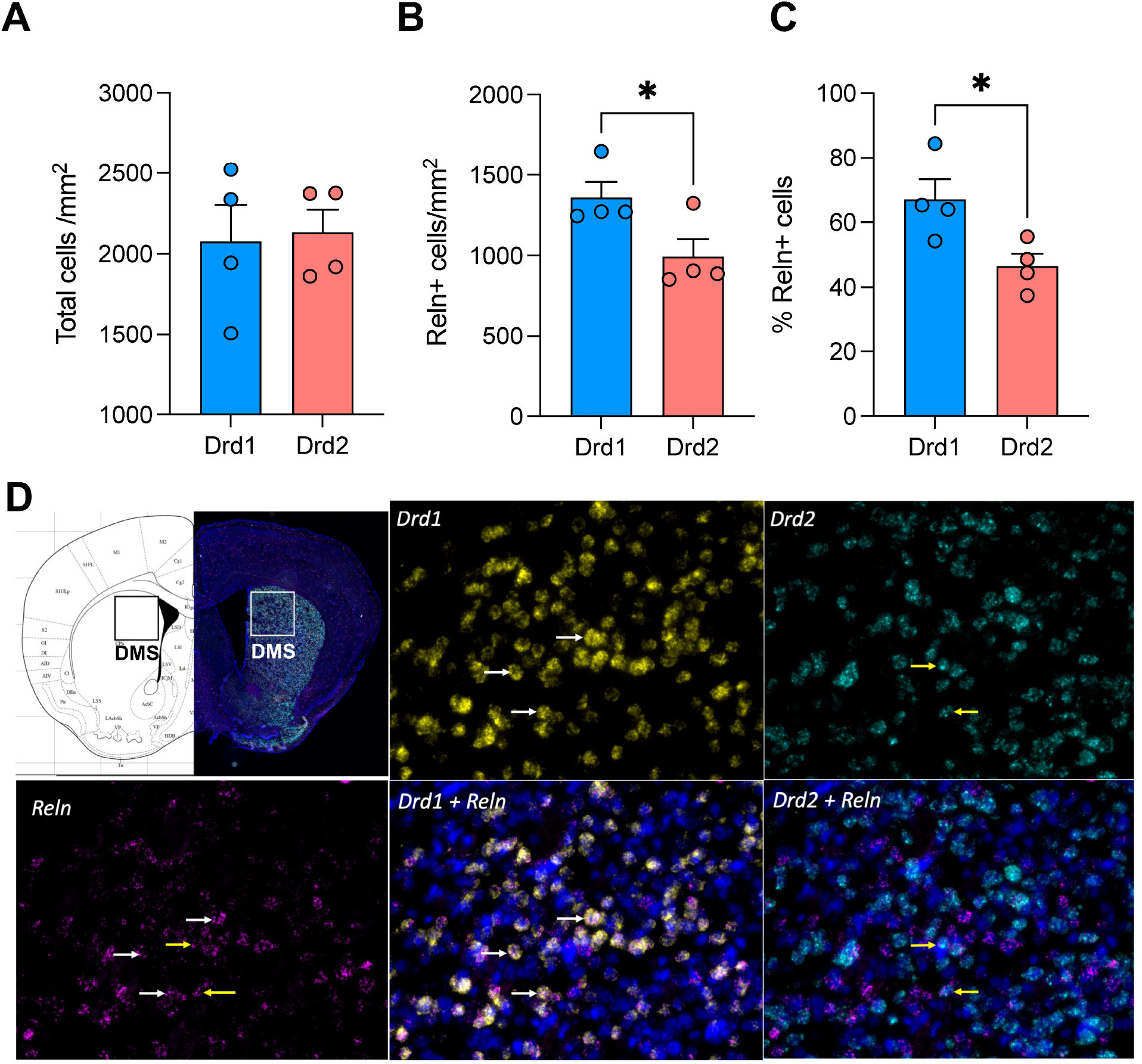
Reelin expression in Drd1 and Drd2 DMS neurons. **A**) Total number of Drd1+ and Drd2+ cells in the DMS. **B**) Total number of Drd1+/Reln+ and Drd2+/Reln+ cells in the DMS. **C**) percent of colocalization Drd1+/Reln+ and Drd2+/Reln+ in the DMS. * *p* < 0.05 vs Drd2. **D**) Representative images for the RNA fluorescent in situ hybridization (FISH).

## Discussion

Our results demonstrated that reduced levels of Reelin influence the locomotor effects of cocaine, but not the cocaine rewarding effects. In our experiments, HR mice showed similar baseline locomotor activity compared to WT, indicating that Reelin deficiency does not influence novelty-induced hyperlocomotion or habituation to the novel environment (Figures 1 and 3). However, when injected with a single dose of cocaine, HR mice showed increased cocaine-induced hyper-locomotion compared to WT. This effect was not due to different sensitivity to cocaine, as demonstrated by the dose-response curve for cocaine-induced locomotor activation shown in Figure 2. The effects on locomotion were confirmed in the cocaine sensitization model, in which HR mice showed higher sensitization to repeated cocaine injections than their WT littermates (Fig. 3). This increased sensitization was also observed after a week of withdrawal (Fig. 3).

These data align with a previous report showing an inhibitory effect of Reelin overexpression on cocaine-induced hyperlocomotion and behavioral sensitization ^35^ confirming a possible role of Reelin signaling in the modulation of the psychostimulant effects of cocaine.

Importantly, we report that the differences observed in cocaine-induced hyperlocomotion and cocaine sensitization did not correlate with differences in the rewarding effects of cocaine. In fact, HR and WT rats developed a similar preference for the cocaine-paired side when subjected to a cocaine place conditioning paradigm. This is also in line with previous data on Reelin overexpressing mice showing no differences in cocaine self-administration rates ^35^ compared to WT mice, demonstrating that manipulation of Reelin does not affect cocaine reward. Considerable evidence suggests that in humans there are sex differences in the pharmacological effects of cocaine and the prevalence of cocaine use disorders. Research in both humans and rodents suggests that women may be more vulnerable to the reinforcing (rewarding) effects of stimulants, with females acquiring cocaine abuse faster and at higher levels than males ^36–39^. Notably, the HR mice exhibit sex differences in the context of neurotransmitter signaling and receptor function ^40,41^. Similarly, HR mice showed sex differences in behavioral disinhibition and stress reactivity following adolescent exposure to THC ^22^. However, our study did not find specific sex-biased responses to cocaine in HR mice, demonstrating that sex does not play a role in the effects of enhanced locomotor activation after cocaine injection in HR mice.

The HR mice showed an increased number of Fos+ cells in the DMS after a single cocaine injection compared to WT mice (Fig. 5). Previous work has shown that cocaine increased Fos expression in the DMS compared to saline, but not in the dorsolateral and ventral striatum ^42^. The increase of Fos positive cells is probably a result of the cocaine-induced increase of dopamine levels in the striatum ^43^, which is hypothesized to increase the activity of MSNs through the activation of dopamine 1 (D1R) and D2 receptors ^44,45^. In particular, MSNs expressing D1R appear responsible for the psychomotor effects of cocaine since mice lacking D1R fail to show the motor-stimulating effects of cocaine ^46^. We found increased expression of Reelin in the DRD1 cells compared to the DRD2 cells in the DMS (Fig. 6). We speculate that Reelin acts as a modulator of DR1 receptor function, which could explain the increased locomotor and DMS neural activations observed after acute injection of cocaine in mice with Reelin deficiency. This hypothesis is reinforced by previous findings demonstrating that both D1 and D2 classes of dopamine receptors are reduced in the striatum of Reeler mice ^47^. Future studies investigating the effects of the manipulation of Reelin expression specifically in DRD1 cells of the DMS will be necessary to test this hypothesis.

The rewarding effects of cocaine are driven by its ability to increase levels of dopamine in the nucleus accumbens ^48,49^. This is reinforced by the fact that blockade of dopamine transmission reduces the rewarding effects of psychostimulants ^50^. In particular, all drugs of abuse are thought to activate the shell subregion of the NAc ^51,52^. In line with the behavioral data that showed no differences in the rewarding effects of cocaine, we did not find any differences in Fos activation in the nucleus accumbens shell of HR and WT mice (Fig 5) after acute cocaine treatment.

## Conclusions

In conclusion, our study indicates that Reelin might have a role in modulating the hyperlocomotory effects of cocaine, but not cocaine reward. Future studies are needed to elucidate the neurobiological mechanisms underlying these findings, such as the relationship between Reelin and the dopaminergic system in the DMS.

## Acknowledgments

This work was supported by the National Institute on Drug Use, USA [DP1DA042232 to FT] and the Brain & Behavior Research Foundation [2020 NARSAD Young Investigator Award to GdG] We thank Havilah Taylor for technical assistance and Dr. Amanda Roberts for the constructive discussions about the experimental behavioral design.

FT designed and coordinated the study. GdG was responsible for the overall behavioral and statistical analysis, data interpretation, figure preparation, and manuscript writing with input from F.T. A.I conducted the Fos IHC experimental design and collected IHC imaging data. A.T. conducted IHC Fos counting with input from A.I. A.N. performed the RNAscope experiment and collected the imaging data with input from A.T. P.M.P performed genotyping.

